# From Ions to Chaos: exploring a whole-brain modelling framework in the mouse

**DOI:** 10.64898/2025.12.04.691835

**Authors:** Simon Adriano Munoz Lagunas, Jeehye An, Leon Stefanovski, Petra Ritter

## Abstract

Computational neuroscience offers powerful tools for understanding brain function in both healthy and diseased states. However, detailed data on microscale mechanisms—such as those involved in genetics and pharmacology—are often derived from mouse experiments and further lack integration into whole-brain models. In this work, we bridge microscale ion-channel dynamics and macroscale network behavior by employing the Larter-Breakspear neural mass model on The Virtual Brain platform with a mouse brain connectome. Our simulations reproduced key findings from prior computational studies with the Larter-Breakspear model, demonstrating its cross-species translation to the mouse. This included the emergence of chaotic dynamics and network synchrony as a function of coupling and delay parameters. Furthermore, we demonstrate that Calcium, Sodium, and Potassium-related parameters each critically shape the global dynamic regimes and that the response to external stimuli is highly sensitive to calcium concentrations. This work represents the first integration of the Larter-Breakspear model into a mouse-scale connectome, validating previous results in a new context. In addition, it establishes a framework for a detailed exploration of synchrony, ion dependency, and stimulus reactivity. By linking molecular simulations, genetic data, and mouse electrophysiology, this approach holds promise for mechanistic insights into neuropsychiatric disorders, such as schizophrenia and epilepsy.

## 1 Introduction

The field of computational neuroscience has emerged as an interdisciplinary science that employs advanced mathematical models and simulation tools to understand the brain’s dynamics. Thus, such computational frameworks can bridge the gap between cellular-level mechanisms of healthy cognition and disease, as well as global brain activity. It provides insights that span from the microscopical scale, where the behavior of individual neurons and ion channels is paramount, to large-scale network interactions that underlie complex cognitive functions. Crucially, the success of such computational approaches relies on the availability of high-quality and multi-scale empirical data. Data derived from imaging modalities, such as functional MRI or diffusion tensor imaging, may provide the necessary constraints and validation for network-level models that describe the large-scale connectivity and dynamics of brain regions.

One of the significant challenges facing the field is that high-resolution microscopic data are predominantly obtained from rodent models, for example, from genetic knock-out mouse models (Groza et al., 2023). Rodents, particularly mice and rats, have provided a wealth of detailed experimental information thanks to the development of well-tailored electrophysiological and imaging techniques specifically designed for these species. Factors like synaptic transmission dynamics and the maintenance of ion gradients across the neuronal membrane may serve as critical links. Incorporating these micro-level variables into network models enables the simulation and analysis of complex brain states with greater biological realism.

We explore the established Larter-Breakspear model within a mouse connectome framework in this context. The Larter-Breakspear model is particularly well-suited for this endeavor as it incorporates a detailed representation of ion channel characteristics, providing a mechanistic link between microscale electrophysiological properties and macroscale network dynamics. By implementing this model on a structural connectivity network derived from mouse data, we aim to assess how well integrating such detailed microscale mechanisms can explain and predict large-scale brain phenomena.

While Breakspear et al. (2003) and Breakspear and Stam (2005) described the dynamics of the Larter-Breakspear system, and Chesebro et al. (2023) conducted an initial bifurcation analysis of its dynamic behavior, the application of this system to a biophysically realistic connectome remains unexplored. To investigate the chaotic behavior of the Larter-Breakspear model within a weighted connectome environment that incorporates distance-dependent time lags, we utilized The Virtual Brain (TVB; http://www.thevirtualbrain.org/, Schirner et al. (2022); Sanz-Leon et al. (2015); Ritter et al. (2013)).

A significant advantage of this approach is its use of weighted connectomes and time lags based on measured and averaged tract lengths in neuronal circuits. It provides TVB with a robust biophysical foundation, unlike models that employ binary connectomes and uniform time lags across regions. The intrinsic chaotic nature of the Larter-Breakspear system is further accentuated when modeling interconnected neuronal populations, resulting in fascinating chaotic dynamics. Notably, two distinct chaotic regimes emerge—chaotic oscillations and chaotic n-cycles—that resemble their non-chaotic counterparts in a single population system but are fundamentally chaotic.

Previous studies have examined the emergent dynamics produced by the Larter-Breakspear model, focusing on factors beyond its chaotic nature. Breakspear et al. (2003) demonstrated that strong coupling results in near-global synchronization, while weaker coupling leads to transitions between phases of synchronization and desynchronization. Moderate coupling values create synchronized clusters. However, they did not account for the effects of conduction delays. Heitmann and Breakspear (2018) built upon this work by introducing constant conduction delays across all nodes and using a binary structural connectome from the macaque brain. They found that strong coupling, combined with conduction delays ranging from 6 to 10 ms—corresponding to the fast rhythm of the Larter-Breakspear system—produced distinct synchronized clusters. In contrast, weaker coupling and/or shorter delays resulted in more disorganized clusters that experienced periods of disorder and reorganization. Shorter delays increased tendencies for global synchronization, though they were often interspersed between states of disorder.

Our investigation aims to validate the model’s ability to simulate neuronal activity patterns that are both biophysically realistic and consistent with experimental observations. This approach holds promise for future translational research. We will focus on reproducing results previously obtained with the Larter-Breakspear model in both artificial and human networks to establish a cross-species perspective. Additionally, we will demonstrate how simulation parameters influence the system’s synchrony and examine how the ions Calcium, Sodium, and Potassium affect the overall dynamics and their sensitivity to external stimuli. We offer a novel application of a detailed and biophysically plausible brain network model in mice, which will facilitate translational research based on these rodent models.

## 2 Results

The results of our investigation into the application of the Larter-Breakspear model to mouse data are presented in five main parts. We first show (1) the fitting of the model with mouse empirical data, then present (2) an exploration of chaotic network dynamics in the fitted model, highlighting different methods of assessing chaoticity in complex, coupled systems. In the following sections, we demonstrate how specific model parameters shape network behavior. Specifically, we demonstrate (3) how coupling parameters and conduction delay shape network synchronization patterns, (4) how varying ion channel parameters lead to different behavioral regimes, and finally, (5) how calcium levels modulate the simulated system’s responsiveness to external stimulation.

### 2.1 Reproduction of Empirical Functional Connectivity Data from Mice

As a first step, we implemented the Larter-Breakspear model to reproduce Functional Connectivity (FC) data from mice. We used publicly available mouse local field potential (LFP) recordings from the Allen Institute. We used LFP recordings from six mouse brain regions that are also present in the structural connectome (see 4.1.2). A stochastic grid search was then used to optimize model parameters.

Fig. 1B shows the best-fit FC matrix obtained from the stochastic grid search optimization, while Fig. 1A shows the empirical FC matrix. Since the FC matrix is always symmetric, we only display the lower triangle of the matrices. The best-fit parameter combination, identified via optimization of root mean squared error, was revalidated using a Pearson correlation between simulated and empirical FCs. This yielded a correlation of *ρ*(13) = 0.8914, p = 8.134 · 10^−6^. This strong correlation indicates that the Larter-Breakspear system with complex parameter space can effectively reproduce realistic functional connectivity properties.

**Figure 1:**
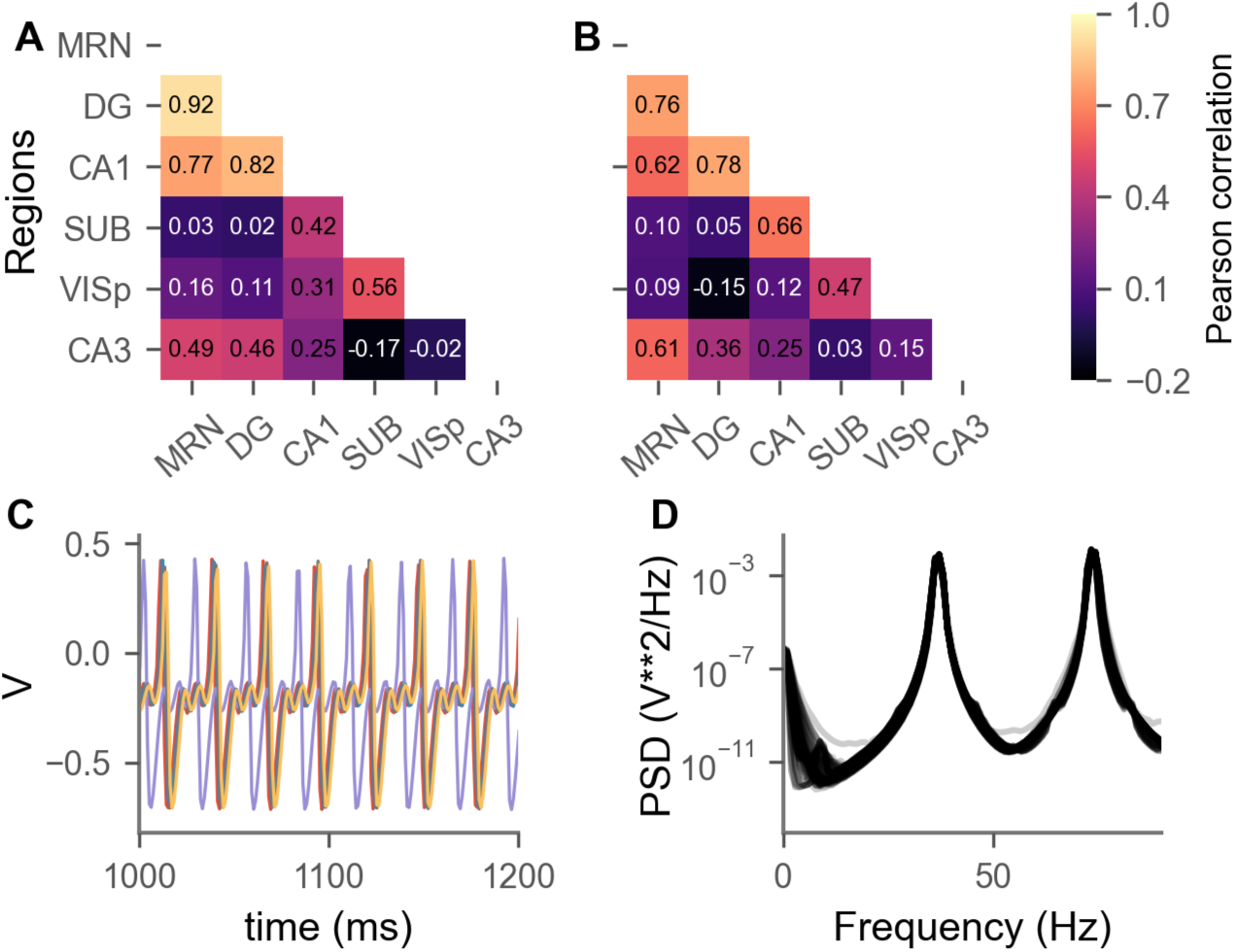
The (A) empirical FC from mouse LFP data and the (B) winning FC matrix from stochastic grid search optimization. The empirical and simulated FCs have a Pearson correlation of *ρ* = 0.8914 and RMSE = 0.132. (C) Example time-series from 5 randomly selected nodes from the fitted Larter-Breakspear network system. (D) PSD of the best-fit Larter-Breakspear network system. MRN=midbrain reticular nucleus, DG=dentate gyrus, CA1=field CA1, SUB=subiculum, VISp=primary visual area, CA3=field CA3.

Fitting the FC of a neuronal system to empirical data can have the risk of obtaining different dynamics than are usual, since the dynamical regimes are not part of the optimization. The dynamics of 5 different nodes of the fitted Larter-Breakspear model in Fig. 1C show that the dynamics are in fact the chaotic non-repeating oscillations that are typical for the Larter-Breakspear neuronal population system, which are described in Breakspear et al. (2003), Chesebro et al. (2023) in single population models and which we further analyse in 2.2. These chaotic n-cycle oscillations are a mix two timescales: a fast one with smaller oscillations and a slow one with higher peaks and a refractory period due to the inhibitory feedback. The power spectral density (PSD) plot (Fig. 1D) indicates two peaks in power, at around 37 Hz and at around 73 Hz. The first peak corresponds to the slow rhythm of the model that creates the big ‘spikes’, led by inhibitory feedback. The second peak, being a multiple of the first, is a result of harmonics.

### 2.2 Exploring Chaotic Network Dynamics

The Larter–Breakspear model encompasses a broad spectrum of dynamical behaviors, including fixed points, periodic oscillations, and complex chaotic regimes. A comprehensive understanding of these dynamics is fundamental to its effective application in modeling neuronal population activity. When extended to interacting populations, the model exhibits an amplification of chaotic dynamics, thereby providing a robust framework for the investigation of emergent phenomena in large-scale neuronal networks.

Extending the analysis by Breakspear et al. (2003) that used single population models, we used the variance of the excitatory threshold (*δ_V_*) as a bifurcation parameter in our network models (Fig. 2). This revealed varying emergent dynamics, consisting of chaotic oscillations and chaotic n-cycle regimes at different frequencies. In the chaotic oscillatory regime (Fig. 2A), the excitatory current is characterized by fast, small-amplitude oscillations that vary in amplitude. As illustrated in Fig. 2B, the chaotic n-cycle regime is characterized by unpredictable, smaller oscillations preceding large spikes. Additionally, the change between small and large oscillations occurs deterministically without needing additional noise. Once *δ_V_* exceeds a certain threshold, the system consistently exhibits the classical aperiodic chaotic oscillations described in Breakspear et al. (2003). In addition, increasing *δ_V_* resulted in higher frequency dynamics. Lastly, we see that these chaotic simulated systems remain bounded to their attractor and do not diverge indefinitely.

**Figure 2:**
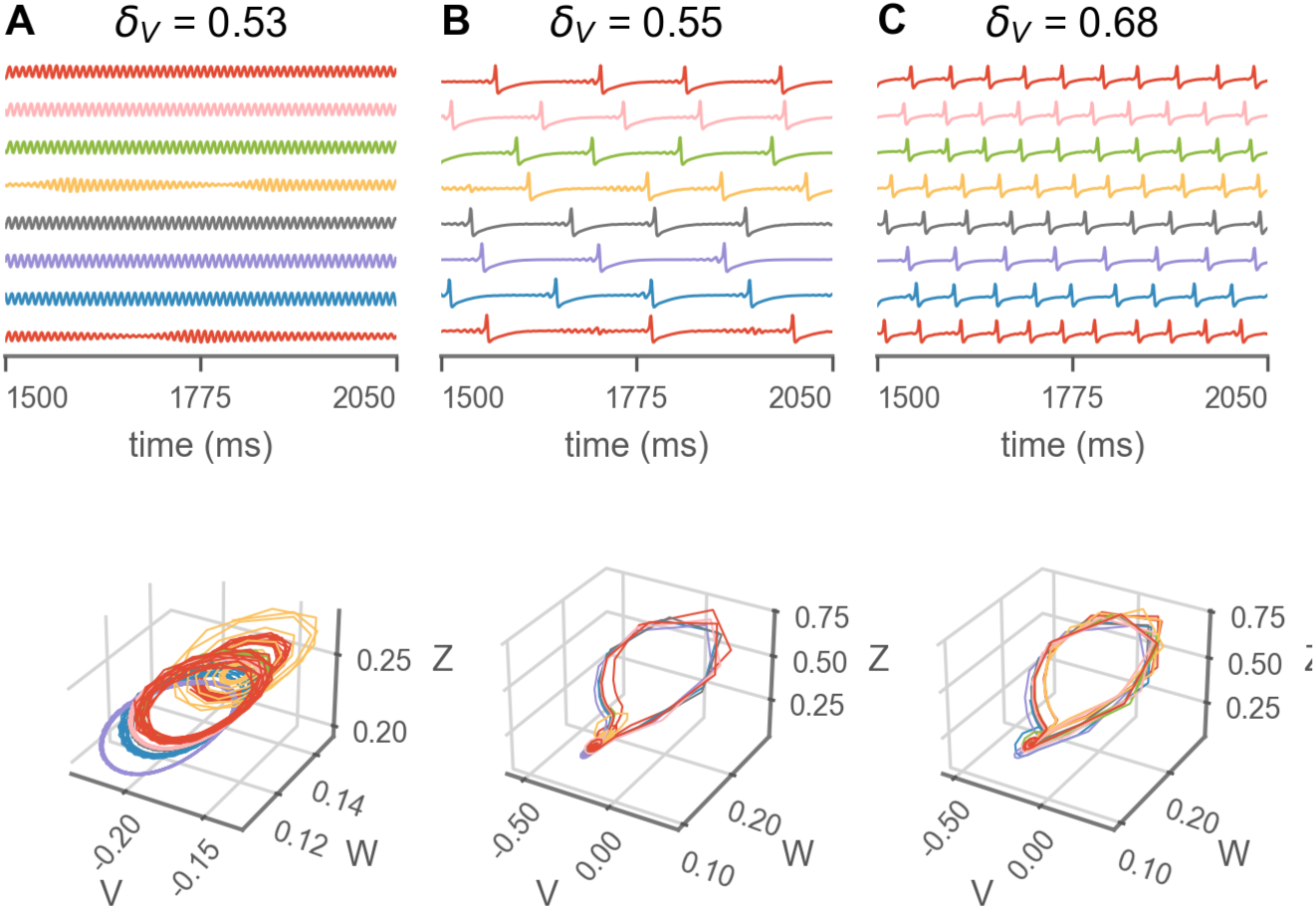
Time series of 8 randomly chosen nodes from simulation using the Larter-Breakspear model with mouse connectome using default parameter values except for varying *δ_V_*. Changing values of *δ_V_* leads to different emergent dynamics. Other parameters were set as the default from Sanz-Leon et al. (2015) in Table 1. The top row shows time-series of the excitatory populations and the bottom row depicts the corresponding phase plane.

**Table 1:**
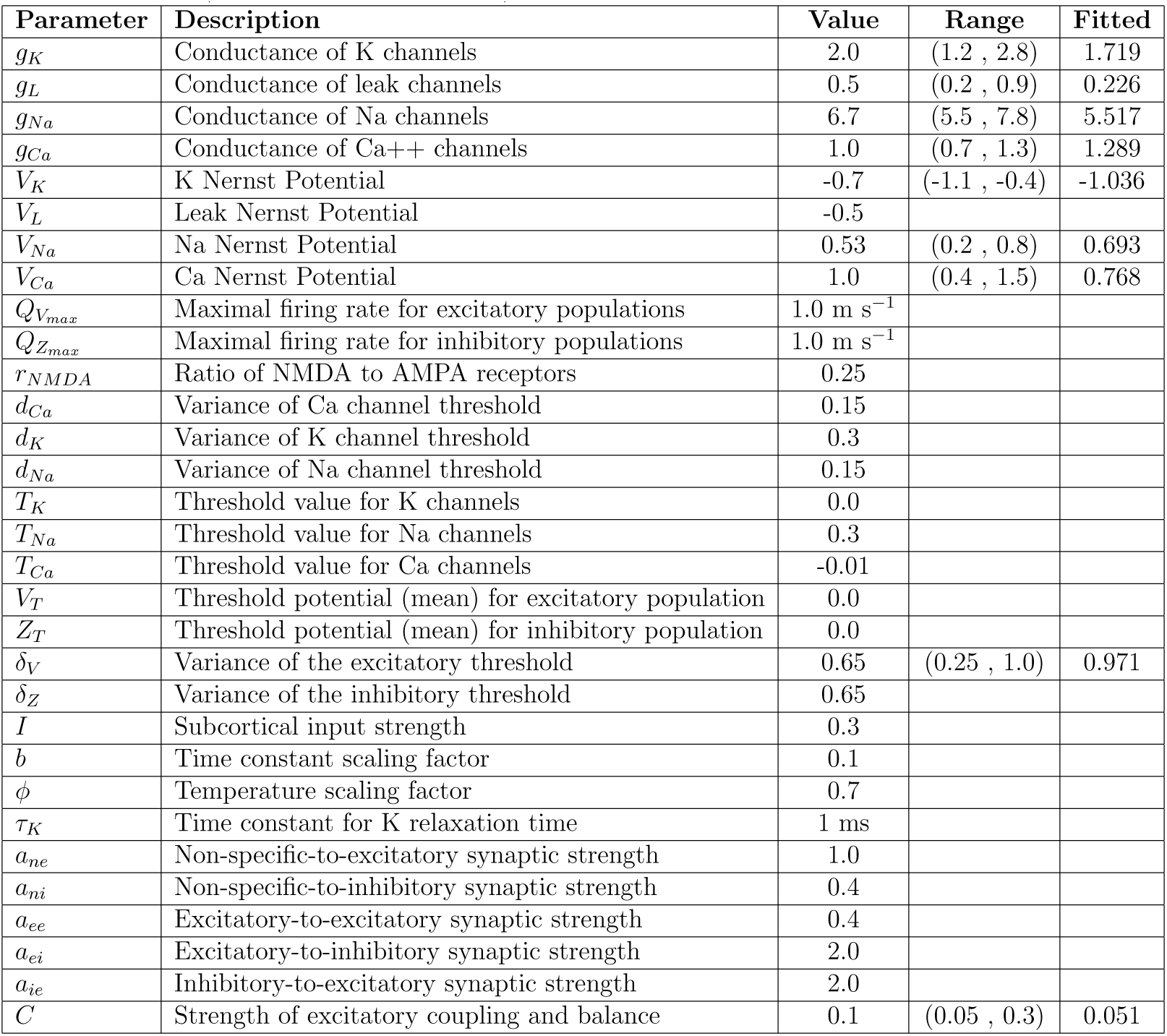
Parameters of the Larter-Breakspear Model taken from Sanz-Leon et al. (2015). *a_ee_* is set to default value from TVB. Parameter ranges and fitted values (rounded to 3 digits) for the stochastic grid search optimization.

| Parameter | Description | Value | Range | Fitted |
| --- | --- | --- | --- | --- |
| $g_K$ | Conductance of K channels | 2.0 | (1.2 , 2.8) | 1.719 |
| $g_L$ | Conductance of leak channels | 0.5 | (0.2 , 0.9) | 0.226 |
| $g_{Na}$ | Conductance of Na channels | 6.7 | (5.5 , 7.8) | 5.517 |
| $g_{Ca}$ | Conductance of Ca++ channels | 1.0 | (0.7 , 1.3) | 1.289 |
| $V_K$ | K Nernst Potential | -0.7 | (-1.1 , -0.4) | -1.036 |
| $V_L$ | Leak Nernst Potential | -0.5 | | |
| $V_{Na}$ | Na Nernst Potential | 0.53 | (0.2 , 0.8) | 0.693 |
| $V_{Ca}$ | Ca Nernst Potential | 1.0 | (0.4 , 1.5) | 0.768 |
| $Q_{V_{max}}$ | Maximal firing rate for excitatory populations | 1.0 m s<sup>-1</sup> | | |
| $Q_{Z_{max}}$ | Maximal firing rate for inhibitory populations | 1.0 m s<sup>-1</sup> | | |
| $r_{NMDA}$ | Ratio of NMDA to AMPA receptors | 0.25 | | |
| $d_{Ca}$ | Variance of Ca channel threshold | 0.15 | | |
| $d_K$ | Variance of K channel threshold | 0.3 | | |
| $d_{Na}$ | Variance of Na channel threshold | 0.15 | | |
| $T_K$ | Threshold value for K channels | 0.0 | | |
| $T_{Na}$ | Threshold value for Na channels | 0.3 | | |
| $T_{Ca}$ | Threshold value for Ca channels | -0.01 | | |
| $V_T$ | Threshold potential (mean) for excitatory population | 0.0 | | |
| $Z_T$ | Threshold potential (mean) for inhibitory population | 0.0 | | |
| $\delta_V$ | Variance of the excitatory threshold | 0.65 | (0.25 , 1.0) | 0.971 |
| $\delta_Z$ | Variance of the inhibitory threshold | 0.65 | | |
| $I$ | Subcortical input strength | 0.3 | | |
| $b$ | Time constant scaling factor | 0.1 | | |
| $\phi$ | Temperature scaling factor | 0.7 | | |
| $\tau_K$ | Time constant for K relaxation time | 1 ms | | |
| $a_{ne}$ | Non-specific-to-excitatory synaptic strength | 1.0 | | |
| $a_{ni}$ | Non-specific-to-inhibitory synaptic strength | 0.4 | | |
| $a_{ee}$ | Excitatory-to-excitatory synaptic strength | 0.4 | | |
| $a_{ei}$ | Excitatory-to-inhibitory synaptic strength | 2.0 | | |
| $a_{ie}$ | Inhibitory-to-excitatory synaptic strength | 2.0 | | |
| $C$ | Strength of excitatory coupling and balance | 0.1 | (0.05 , 0.3) | 0.051 |

#### 2.2.1 Assessing Chaoticity using Maximal Lyapunov Exponents

To assess whether a deterministic dynamical system is chaotic, one can examine the rate at which initially close trajectories diverge. This rate is quantified by Lyapunchaoticityts, where a positive value indicates chaotic dynamics. The maximal Lyapunov exponent (MLE) is the largest Lyapunov exponent of a system, and it’s often used as a key indicator of chaos in the system. In the single-population Larter-Breakspear system, Breakspear et al. (2003) estimated a positive maximal Lyapunov exponent *λ*^∗^ = 0.015 using a Gramm-Schmidt algorithm. We have extended this finding to the complex chaotic dynamics of the coupled Larter-Breakspear network system. We simulated the Larter-Breakspear network with parameters from Fig. 2C and determined the MLE of all simulated time series using Eckmann et al. (1986)’s method. The MLEs for all analyzed signals were positive, confirming that all populations are chaotic (Fig. 3). We confirmed the positive MLEs with the algorithm from Rosenstein et al. (1993), reinforcing the chaotic nature of the dynamics. While the exact values of the exponents depend on the time grid and algorithm used, the overall conclusion regarding the model’s chaoticity remains consistent.

**Figure 3:**
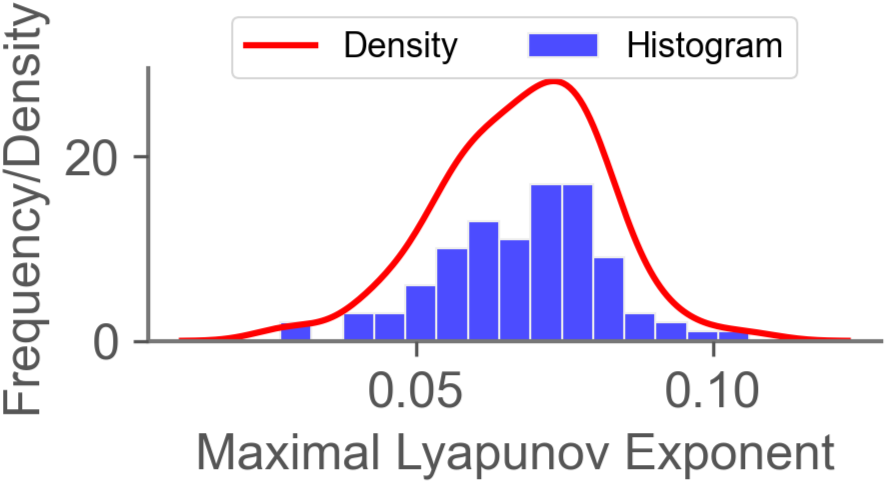
Histogram and density plot of the maximal Lyapunov exponents from all simulated populations in the chaotic n-cycle regime. MLE was determined from the state variable V (excitatory current, 1) using Eckmann et al. (1986)’s method. All values are positive, indicating chaotic dynamics.

#### 2.2.2 Assessing Chaoticity with Poincaré Analysis

As a second way to assess the chaoticity of dynamics, we used Poincaré maps. The Poincaré map is a projection of the system’s dynamics onto a lower-dimensional subspace. Chesebro et al. (2023) used a Poincaré map on the single population Larter-Breakspear model to demonstrate chaos. For a thorough introduction to Poincaré maps and their applications in chaotic systems, see Strogatz (1994). Since there is no one unique way of constructing Poincaré maps, it is the researcher’s task to find a suitable lower-dimensional representation. We present two methods for constructing Poincaré maps, demonstrating the chaotic nature of the dynamics and enabling easy distinction between different chaotic regimes in the lower-dimensional representation.

The first method for constructing Poincaré maps from time series data involves using a planar cut through the 3-dimensional phase space (Fig. 4). The dense clustering of points in the Poincaré map (Fig. 4B, D) indicates frequent, non-repeating plane crossings, revealing the chaoticity of the Larter-Breakspear system.

**Figure 4:**
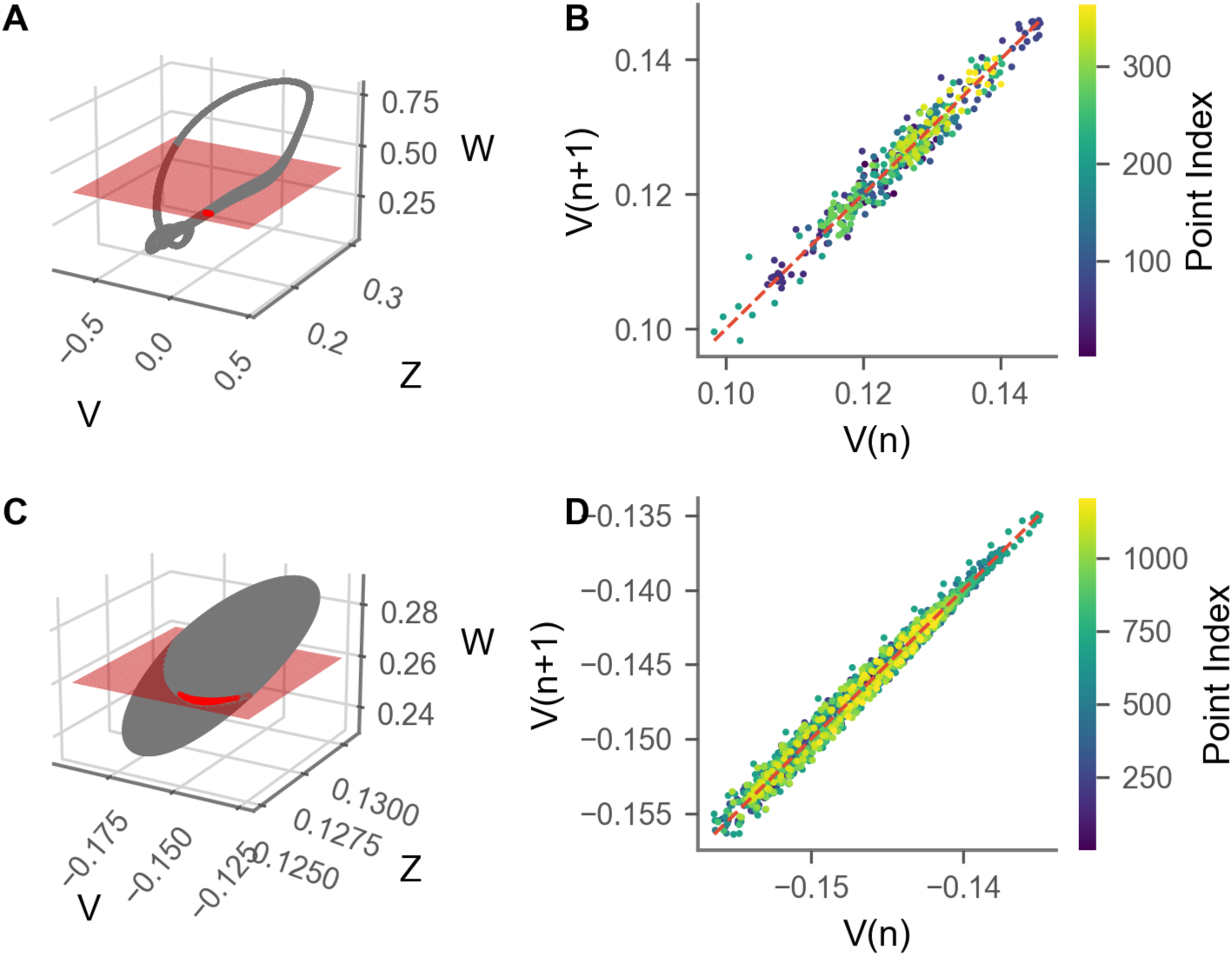
3-dimensional phase space plot and Poincaré map of one population computed with the plane crossing method. (A) and (B) used a fitted model, while (C) and (D) used time series in the oscillatory regime. The heat map of the Poincaré plots shows the crossing index, and the red dashed line indicates the identity line.

Fig. 4A shows the time series of one node of the simulation in the chaotic n-cycle regime, using the best-fit parameters from 2.1 and the Poincaré section surface (red) used to determine Poincaré points. Fig. 4B shows the resulting Poincaré map for a single population in the chaotic n-cycle regime using the plane crossing method. Since we are interested in the dynamics of the excitatory population, the Poincaré map shows the values of the excitatory variable *V* (*n*)*, V* (*n* + 1) at consecutive Poincaré points *n* = 1*, …, N*. As the length of the analyzed time series increases, the set of distinct points in the resulting Poincaré map grows linearly. Importantly, these points remain non-overlapping, indicating that the state space is continuously being explored without revisiting identical state transitions. This observation is illustrated in Fig. 4B,D, where the number of plotted points in the Poincaré map scales with the duration of the simulation, yet no discernible fixed patterns or recurrent loops emerge. A key indicator of chaotic dynamics in Poincaré analysis is the absence of convergence to the identity line. If the system were to converge to a stable fixed point or periodic orbit (as expected in non-chaotic systems), all points would lie on the identity line *V_n_*_+1_ = *V* (*n*), eventually forming a single point or a closed loop. In our case, the map remains diffuse, with no accumulation of points along the identity line, even after long simulation lengths. This strongly suggests the absence of an attractor in the form of a limit cycle or fixed point. Furthermore, the associated heat map visualization in Fig. 4B shows no spatially coherent structures or clustering patterns. Results were consistent with integration time steps of 1 ms and 0.01 ms.

We repeated the analysis on time series in the oscillatory regime (Fig. 4C) and observed similar results as in the analysis with the fitted model. The resulting map in Fig. 4D forms a complex, non-repeating point cloud which centers around the identity line (red dashed line) without collapsing onto it. The heat map of crossing indices reveals no discernible structure, indicating chaotic dynamics.

Secondly, we constructed Poincaré maps using local maximum approach. Fig. 5A shows a snapshot of the excitatory current dynamics from the Larter-Breakspear network model in the chaotic n-cycle regime and its local maxima. The time series reveals smaller local maxima followed by rapid transitions to higher maxima, reflected in the three distinct point clouds on the Poincaré map. Fig. 5B displays the corresponding Poincaré map, which highlights consecutive local maxima, clearly illustrating the system’s chaotic nature. The map’s numerous unique points, depending on the time series length, confirm the chaotic dynamics.

**Figure 5:**
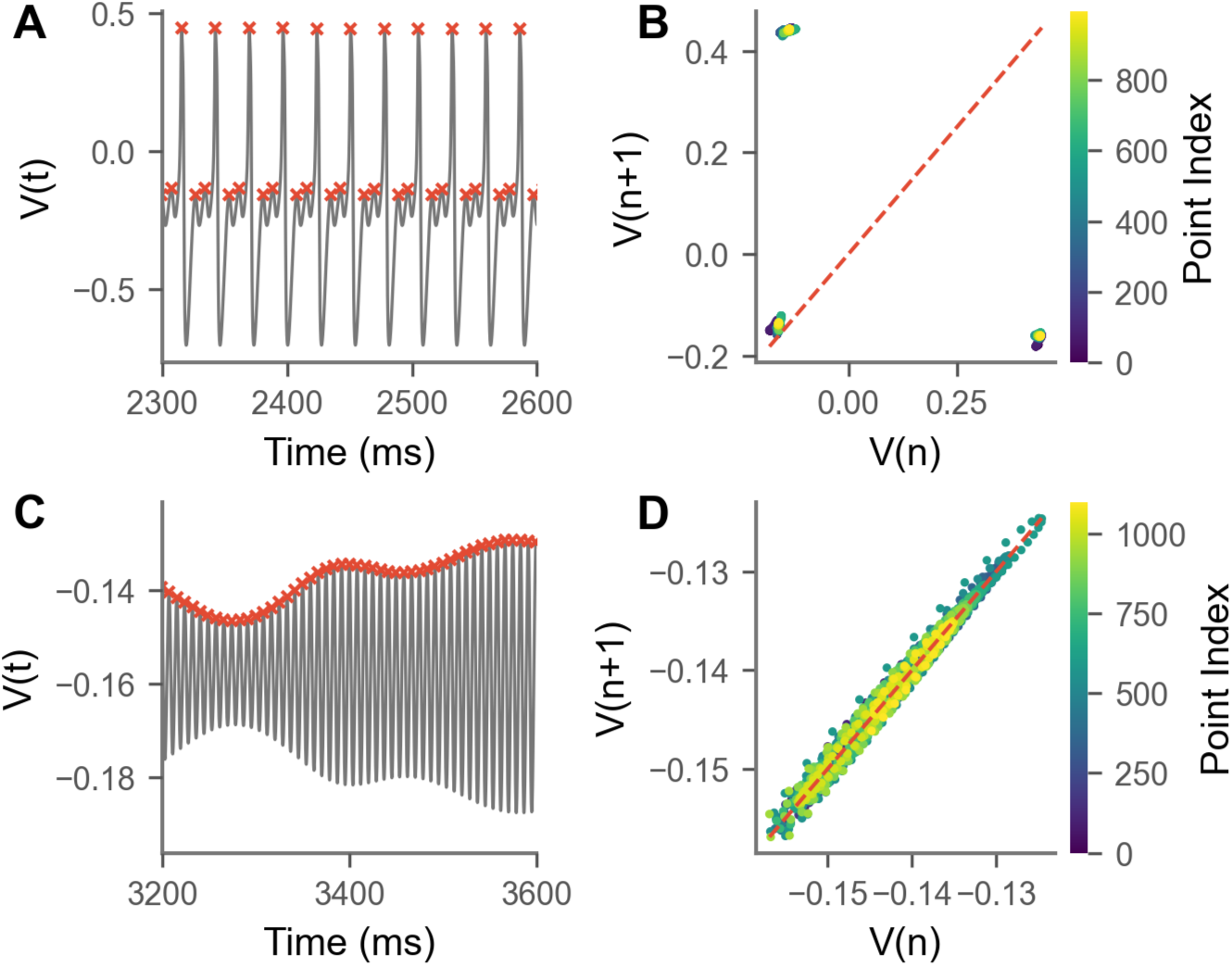
Identification of local maxima, indicated as red crosses, in the excitatory current time series *V* (*t*) of (A) the fitted model, and (C) the chaotic oscillatory regime. (B) Resulting Poincaré map in *V* (*n*)*, V* (*n*+1) axes showing Poincaré points of the excitatorythe current derived through local maximum approach for fitted model and (D) chaotic oscillatory regime. Red dashed line indicates the identity line. The Poincaré points *V* (*n*)*, n* ∈ {1*, …, N* } are ordered by increasing time while the heatmap shows the discrete index *n* ∈ {1*, …, N* } of the Poincaré points.

Fig. 5C shows a snapshot of the time series of the excitatory current and highlights that consecutive local maxima are close but never identical, further emphasizing the system’s chaotic nature. When applying the local maximum method to the excitatory current time series in the chaotic oscillatory regime, the resulting Poincaré map (Fig. 5D) aligns with the identity line, resembling the map obtained via the plane crossing method.

### 2.3 Network Synchronization depends on Coupling and Conduction Delays

We compared network synchronization patterns under two types of conduction delay: constant delays across all nodal pairs (Fig. 6, left column), and distance-dependent heterogeneeous delays (right column). Constant delays at different values produced distinct synchronized clusters or near-global synchronization (e.g. panels A and D, left). In contrast, introducing distance-dependent delays yielded strikingly different patterns, even when average delays were matched between conditions.

**Figure 6:**
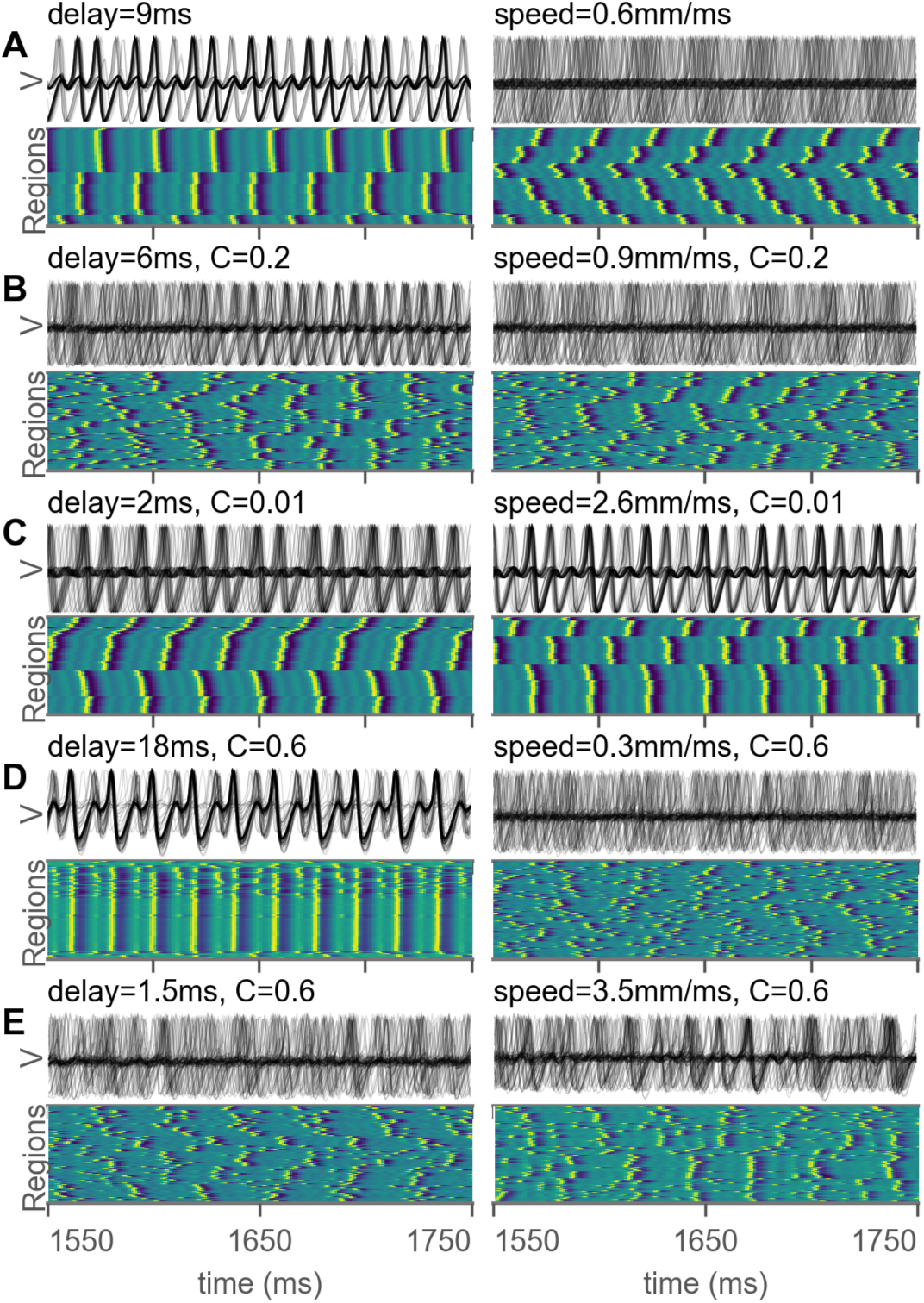
Comparison of emergent dynamics under two different treatments of conduction delay: constant delays (left column) and distance-dependent heterogeneous delays (right column). Conduction speeds for the heterogeneous delay condition were selected to match the average delay of the constant-delay condition to its left. Within each row (A-E), the left and right plots share identical parameters except for the treatment of conduction delays. Top row plots the excitatory current (V), of simulated time-series and the bottom row plots its heatmap, with nodes ordered based on hierarchical clustering.

Previous studies have shown that constant delays that match the model’s intrinsic rhythm led to distinct synchronized clusters, while weaker coupling and/or shorter delays produced more disorganized dynamics (Heitmann and Breakspear, 2018). Meanwhile, short constant delays led to instances of global synchrony interspersed among chaos. Here, we extend the analysis by implementing more realistic brain connectivity by using a weighted structural connectome (Oh et al., 2014) and introducing distant-dependent conduction delays. Conduction delays in TVB are calculated as the estimated tract length divided by a specified conduction speed, so that nodes that are closer to each other communicate faster and vice versa.

Using a constant conduction delay of 9 ms, which matches the fast rhythm of our fitted model, produced synchronized clusters (Fig. 6A, left), reproducing Heitmann and Breakspear (2018). However, these clusters vanished when using distance-dependent delays with matched average delay (Fig. 6A, right). Using a slower delay and/or stronger coupling decreased synchronization tendencies, again aligning with Heitmann and Breakspear (2018). For example, using a shorter constant delay of 6 ms with a stronger coupling of 0.2 (Fig. 6B, left) led to phases of synchronized clusters interspersed by phases of disorder. phases of synchronized clusters interspersed by phases of disorder. However, this observation again vanished when distance-dependent delays were implemented.

When distance-dependent delays are applied, clusters were observed when a combination of relatively faster speeds (leading to shorter delays) and weaker coupling values were applied. An example is shown in Fig. 6C, right column. Synchronized clusters were lost when using slower speeds and/or stronger coupling values.

Strong coupling (*C* = 0.6) combined with a relatively long constant delay led to near-global synchronization (Fig. 6D, left), but this pattern was lost with distance-dependent delays. Instead, we observed disordered, chaotic dynamics without apparent clustering (Fig. 6D, right). Finally, using a strong coupling of 0.6 with either short delay of 1.5 ms or fast conduction speed of 3.5 mm/ms produced chaotic, unpredictable dynamics with brief instances of partial synchronization (Fig. 6E).

In summary, we were able to reproduce previously reported dynamics using constant delays on a weighted mouse connectome. Specifically, a constant delay of 9 ms produced distinct clusters, while shorter delays and/or stronger coupling eliminated or diffused clustering. However, using distance-dependent delays significantly altered these patterns – near-globally synchronized dynamics never emerged, and clustering was weaker or absent and only emerged under specific fast-speed, low-coupling conditions.

### 2.4 Ion Channel Characteristics determine System Dy-namics

The Larter-Breakspear system consists of several parameters, including Nernst potential parameters, that govern calcium, sodium, and potassium ion channel dynamics. We investigated how varying these parameters – *V_Ca_*, *V_Na_*, and *V_K_* – affects network dynamics in our network model of a mouse brain. Recent work by Chesebro et al. (2023) conducted a bifurcation analysis on these parameters in a single population Larter-Breakspear system without conduction delay and showed that changing these parameters changes the dynamical regimes or oscillation frequencies via period-doubling bifurcations. Here, we extend the analysis to our network model and given the complexity of our 98-node network, we focus on exploratory analysis of emergent dynamics rather than a detailed bifurcation analysis.

When *V_Ca_* ≤ 0.45, the system rapidly converges to a stable fixed point (Fig. 7A). As *V_Ca_* increases to around 0.55, most nodes become unstable and begin oscillating around the fixed point with variable amplitudes. Further, increasing *V_Ca_* amplifies the oscillations until a bifurcation occurs, leading to chaotic n-cycle oscillations. At *V_Ca_* of 0.66, spike frequency rises, smaller oscillations decrease, and more populations transition into the chaotic n-cycle regime. By *V_Ca_* ≈ 0.7678, the dynamics become smoother and sharper.

**Figure 7:**
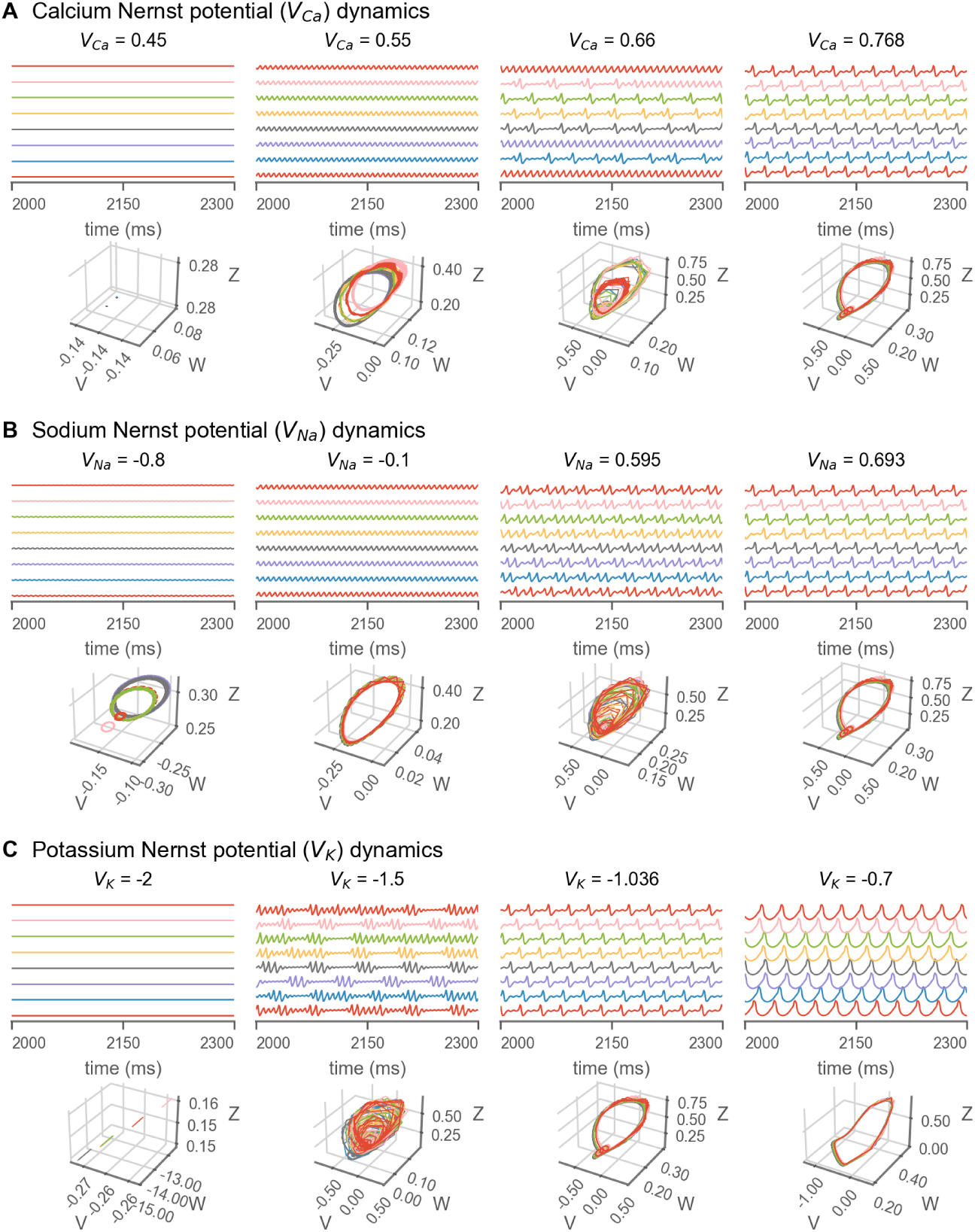
Altering (A) calcium (*V_Ca_*), (B) sodium (*V_Na_*), and (C) potassium (*V_K_*) ion channel Nernst potential parameter values yields different dynamical regimes. In each panel, top row plots the mean membrane potential, *V* (*t*), of 8 randomly selected nodes. The bottom row shows corresponding three-dimensional phase plane of the trajectory of the same nodes. The fitted values of *V_Ca_*, *V_Na_*, and *V_K_* are 0.768, 0.595, and –1.035, respectively.

Changing the sodium Nernst potential (*V_Na_*) produced similar alterations (Fig. 7B). At *V_Na_* ≤ −0.8, the system shows stable fixed points and weak oscillations. At *V_Na_* = −0.1, low-amplitude oscillations appear. Higher values of around 0.595 trigger a transition to chaotic n-cycle oscillations. As *V_Na_* increases to 0.6926 and above, the system stabilizes in the chaotic n-cycle regime with smoother transitions.

For potassium Nernst potential (*V_K_*, Fig. 7C), the system stabilizes at a fixed point at *V_K_* ≤ −2.0. Increasing *V_K_* to around −1.5 introduces non-periodic, chaotic oscillations. At the fitted value of −1.036, the system enters the chaotic n-cycle regime with more regular spikes and smoother transitions. At *V_K_* ≥ −0.7, weak oscillations vanish, and high-amplitude, spiky oscillations dominate.

### 2.5 Sensitivity to Stimuli is modulated by Calcium Levels

The system’s sensitivity to external stimulation varied across dynamical regimes, which were produced by changing calcium Nernst potential parameter. We varied the calcium Nernst potential — as in Fig. 7A — to produce four dynamical regimes: fixed-point, stable oscillatory, and mixed/highly chaotic, and chaotic n-cycle dynamics. We then applied either a single pulse or a train of repeated stimuli to the right primary motor area and observed the system’s responses (Fig. 8).

**Figure 8:**
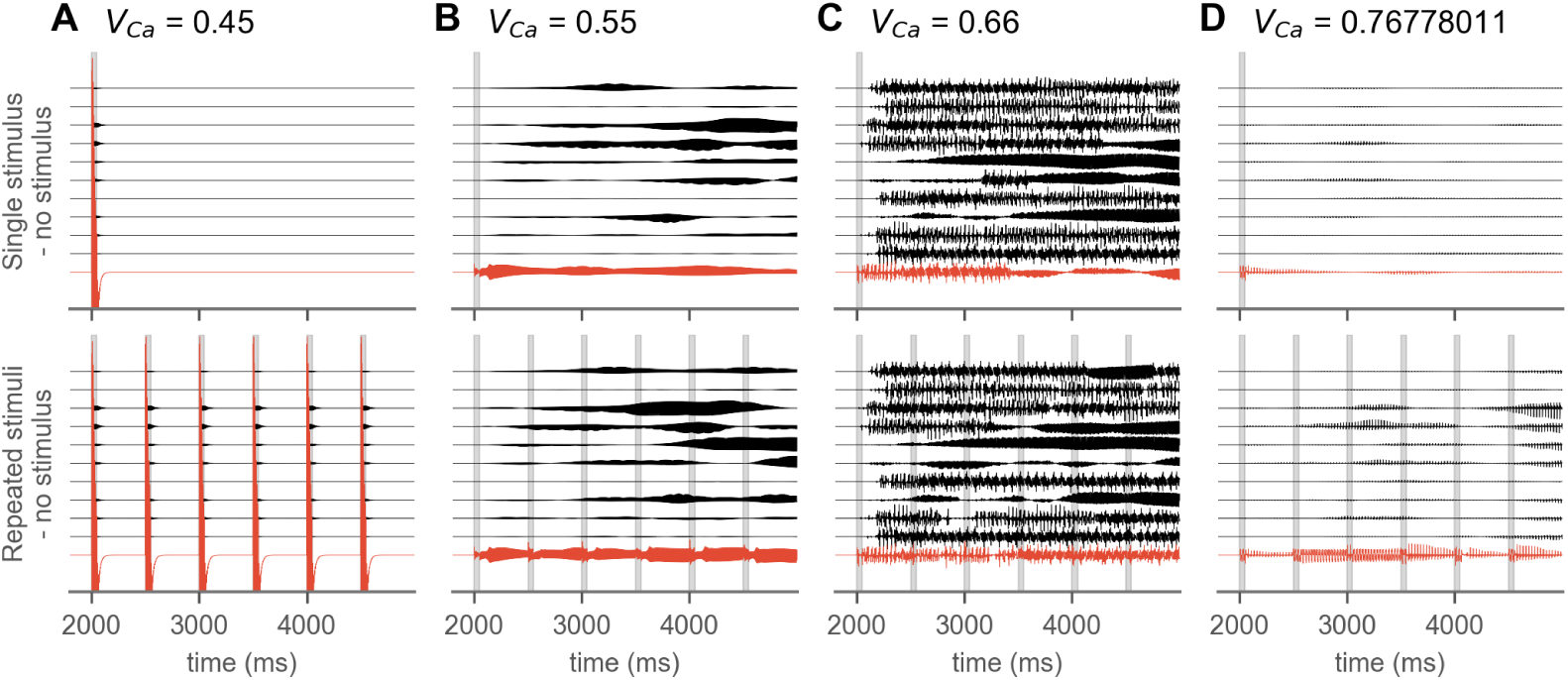
Calcium channel activity modulates the simulated system’s sensitivity to external stimuli. The difference between stimulus and no-stimulus conditions from simulations with identical initial conditions is plotted, thereby isolating the effect of stimulation. Stimuli were delivered to right primary motor area (plotted in red) and were delivered either only once (top row) or repeatedly every 500 ms (bottom row). Ten regions with the strongest structural connectivity to the stimulated region are shown. Stimuli delivery is indicated by gray vertical bars. The different dynamical regimes produced by varying the calcium Nernst potential (*V_Ca_*) are shown in Fig. 7A.

Fig. 8 illustrates the effects of stimulation by plotting differences between stimulated and non-stimulated runs with identical initial conditions. Since the simulations are deterministic (i.e., no noise), these differences effectively isolate the impact of external stimulation. Full plots showing all 98 nodes of the mouse connectome are provided in the supplementary materials (Fig. S5).

In the fixed-point regime (*V_Ca_* = 0.45, Fig. 8A), stimulation triggered immediate but transient disturbances across nodes, without any lasting effects. In contrast, in the stable oscillatory regime (*V_Ca_* = 0.55, Fig. 8B), even a single stimulus induced enduring effects across nodes, and repeated stimulation introduced additional changes over time. In the chaotic n-cycle regime (*V_Ca_* = 0.77, Fig. 8D), a single stimulus had only a small initial impact, but repeated stimulation cumulatively amplified perturbations over time. The system was particularly vulnerable in the highly chaotic mixed regime (*V_Ca_* = 0.66, Fig. 8C), where even a single stimulus caused immediate, widespread, and persistent alterations.

To better understand how stimulation influenced network behavior, we compared PSDs and dynamical functional connectivity (dFC) across the three conditions: No stimulation, single stimulation, and repeated stimulation. Overall, external stimulation primarily altered dFC patterns over time while leaving signal frequencies largely unaffected (Fig. 9). The exception was the fixed-point regime (*V_Ca_* = 0.45, Fig. 9A), where stimulation increased power due to transient activation of previously inactive nodes.

**Figure 9:**
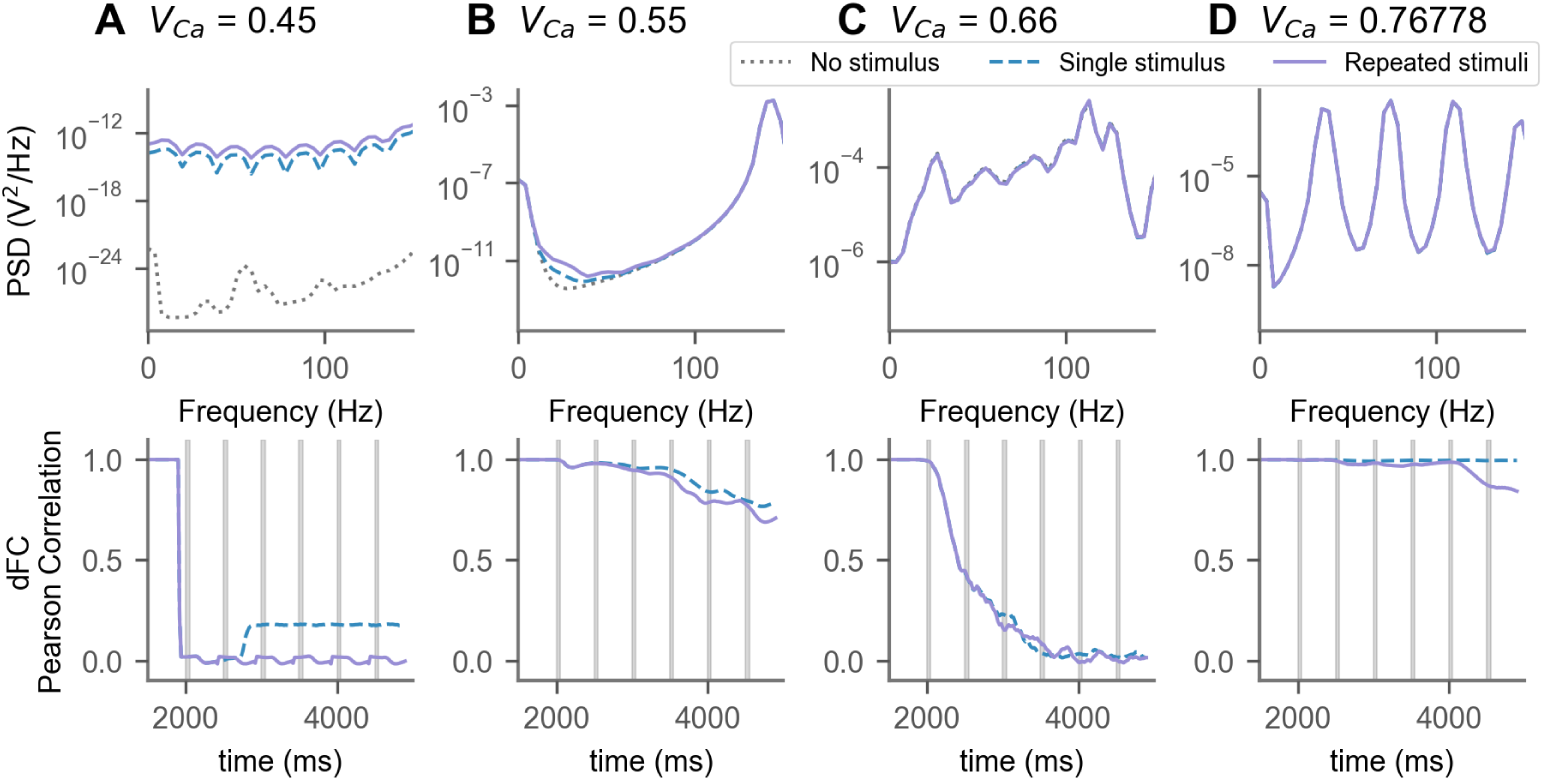
External stimulation primarily affected dFC patterns over time while leaving power spectra essentially unchanged. The top row shows the average PSDs for each stimulation condition. For *V_Ca_* = 0.45 (panel A), overall power without stimulation (gray dotted line) is very low since most nodes are in a fixed-point regime, i.e., without detectable activity. Increased power with stimulation reflects a transient increase in activity induced by the stimuli. For other *V_Ca_* values, PSDs across conditions show no notable differences, indicating that stimulation did not significantly alter signal frequencies. The bottom row shows Pearson correlations between dFC matrices from simulations with and without stimulation. Higher chaoticity is linked to greater perturbability of functional connectivity over time.

In contrast, dFC patterns were strongly influenced by system’s chaoticity and partially by stimulation type. In stable oscillatory systems (*V_Ca_* = 0.55, Fig. 9B), a single stimulus moderately altered dFC over time (dashed blue line), and repeated stimulation led to slightly greater cumulative changes over time (purple solid line). In stable chaotic systems (*V_Ca_* = 0.77, Fig. 9D), dFC remained robust to a single perturbation, but repeated stimulation had the correlation drop to around 0.85 by 5000 ms.

Highly chaotic systems (*V_Ca_* = 0.66, Fig. 9C) were extremely sensitive to external input; Even a single stimulus rapidly drove dFC correlation to near-zero, indicating complete reorganization. A similar drop was observed with the fixed point regime (*V_Ca_* = 0.45, Fig. 9A), although this may reflect activation of a previously silent system rather than network restructuring.

## 3 Discussion

In this work, we demonstrated the potential of whole-brain simulations with the Larter-Breakspear Model in a mouse brain. Our work allows for several applications, ranging from cellular ion concentrations to chaotic phenomena in brain behavior. First, we showed how the Larter-Breakspear model can accurately reproduce mouse brain functional connectivity, achieving a high Pearson correlation (*ρ*(13) = 0.8914) between empirical and simulated FC matrices. Second, we used maximal Lyapunov exponent and Poincaré analysis to examine chaoticity of our complex, coupled system. Then, we reproduced previous observations on synchronization patterns using constant conduction delay and showed how distance-dependent conduction delays significantly alter these patterns. We also used ion channel Nernst potential parameters to place the system in different dynamical states. Finally, we demonstrate that altering the calcium Nernst potential parameter produces systems with different sensitivities to external stimulation.

The first challenge of this work was to reproduce highly complex empirical data obtained from invasive recordings in the mouse brain. Fitting the network model to empirical mouse brain LFP data produced a strong correlation between simulated and empirical FC matrices (*ρ*(13) = 0.8914, p = 8.134 · 10^−6^), suggesting that the model successfully captures key aspects of neuronal dynamics. A notable strength of this approach is the direct use of the Allen Institute’s mouse brain connectome (Oh et al., 2014), which enabled alignment between simulated brain regions and anatomical coordinates of empirical data. This spatial correspondence provides a bridge between the simulated model and biology, strengthening the interpretability of our findings. Nonetheless, the empirical LFP data covered only six regions, while the model simulates 98, limiting the fitting process to a 6 × 6 FC matrix.

The stochastic grid search optimization method, although effective, has limitations due to the ample parameter space and computational demands. A complete brute-force search is unrealistic due to computational demands. However, a parallelized stochastic grid search on a high-performance computing cluster provided reliable results.

However, every approach of data fitting also needs to address the problem of overfitting to avoid artificial, non-translatable results. To evaluate overfitting phenomena, we monitored the RMSE evolution across multiple parallel fitting runs on the HPC cluster, as shown in the supplementary material (Fig. S1). Our methods effectively balanced computational efficiency and fitting accuracy, demonstrating the robustness of the optimization approach. To control for overfitting, we ensured that the stochastic grid search was run for a sufficiently long duration and across a diverse parameter space, allowing the RMSE between empirical and simulated FC matrices to converge consistently across independent runs.

The chaotic nature of the dynamical model offers several benefits but also represents a highly complex approach that requires proper mathematical methods for its effective analysis. The analysis of MLEs calculated from the simulated time series confirmed the chaotic nature of the Larter-Breakspear network dynamics. Consistently positive MLE values for all populations indicate a sensitive dependence on initial conditions, characteristic of chaos. Although MLE estimation from time series data lacks the precision of analytical solutions, results align with earlier single-population model findings (Breakspear et al., 2003), extending evidence of chaotic dynamics to the network model.

Poincaré map analysis proved effective for characterizing chaotic dynamics. The plane crossing method facilitated the visualization of chaotic behavior and served as a valuable diagnostic to confirm the presence of chaos in the system. In our analysis, we employed this method to verify the existence of chaotic trajectories by examining the scattering and recurrence patterns of intersections. However, it required manual selection and positioning of the transversal plane to ensure meaningful projections. In contrast, the local maximum method yielded higher precision by isolating successive local maxima in the time series, thereby minimizing ambiguity in point selection.

This approach proved particularly valuable for distinguishing between different types of chaotic regimes, such as chaotic *n*-cycles and chaotic oscillatory states. The alignment of points along the identity line in the Poincaré map emerged as a key indicator for differentiating these regimes, highlighting the method’s relevance for the study of complex oscillatory dynamics.

Besides chaoticity, synchronization is a paramount phenomenon not only in the biological brain, but also in the system of coupled Larter-Breakspear oscillators. Herein, it is significantly influenced by the choice of conduction delays. When a constant delay was applied across all nodes, and the delay matched the model’s intrinsic rhythm, stable synchronized clustering emerged. This occurs because incoming signals from other nodes arrive in phase with each node’s natural oscillation, and under moderate to low coupling, this leads to the formation of multiple phase-locked states. A similar phenomenon was previously demonstrated using a binary macaque connectome (Heitmann and Breakspear, 2018), and we replicate this finding using a weighted mouse connectome.

We show that introducing distance-dependent delays significantly alters this dynamic. In many cases, synchronized clusters either diffuse or disappear entirely. Instead, clusters were only observed under conditions of faster conduction speeds (shorter delays) and weaker coupling. Heterogeneous conduction delays introduce additional variability into the system, resulting in a broader spectrum of phase relationships between nodes. We propose that faster conduction speeds may help reduce this variability, while weaker coupling may reduce the influence nodes have on each other. Conversely, strong coupling may increase inter-nodal influence and consequently magnify variability.

An exponential relationship between connection length and weight, called the exponential distance rule (EDR), is a well-established feature across the mammalian species (Donahue et al., 2016; Ercsey-Ravasz et al., 2013). Yet, many models overlook this by assuming uniform delays regardless of distance. The decreased tendency to form synchronized clusters under distance-dependent delays implies that intrinsic temporal heterogeneity may serve a functional role — perhaps limiting excessive global synchronization. In the biological brain, conduction delays vary by region and species, influenced by factors such as myelination and axon diameter (Wang et al., 2008). Although our model used a uniform conduction speed as a simplification, we show even this plays a significant role in shaping emergent patterns of synchronization and metastability.

A great advantage of the presented whole-brain modelling approach is the possibility to incorporate data from the subcellular level of ions, as well as the potential to assess its reaction towards external stimuli. In analyzing the effects of ion channel gradients, we show that bifurcations observed in single-node systems (Chesebro et al., 2023) persist in a complex, weighted network model of the mouse brain. For instance, depolarizing calcium channel gradients shifted the system’s dynamics from chaotic oscillations to stable limit cycles, with an intermediate mixed regime. Notably, different regimes exhibited varied sensitivity to external stimulation, underscoring the role of calcium gradients in shaping dynamic behavior.

Our results show that the system’s responsiveness to external stimulation depends strongly on its underlying dynamical regime, which can be tuned through ion channel parameters like calcium Nernst potential. In highly chaotic regimes, even a single stimulus triggered widespread and lasting reorganization of dFC. In contrast, more stable regimes resisted transient perturbations but exhibited cumulative changes when exposed to repeated stimulation.

Interestingly, external stimulation primarily affected dFC patterns over time, while PSDs remained largely unaffected. This suggests that stimulation selectively targets temporal reorganization and transitions between network states rather than directly modulating signal frequencies. These observations highlight the importance of considering the brain’s intrinsic dynamical state when designing neuromodulation protocols or interpreting neural responses to external perturbations. Systems operating near chaotic regimes may exhibit greater plasticity and responsiveness but also carry a greater risk of overstimulation. Conversely, more stable systems may require repeated inputs to achieve meaningful reorganization.

In summary, this study highlights the value of biologically informed network models in simulating realistic brain dynamics and lays the groundwork for future research into how computational models bridge empirical observations and theoretical predictions in neuroscience. Future work should incorporate additional empirical data, covering more brain regions, to refine parameters and further enhance the model’s validity.

## 4 Methods

The following describes the data sources, modeling framework, and analysis pipeline used to link mouse structural connectivity, spontaneous LFP recordings, and biophysical network simulations. Thus, we introduce the tracer-derived mouse connectome and detail the preprocessing of LFP epochs. Further, we outline how these empirical data were used to constrain The Virtual Brain simulations by using the Larter-Breakspear neural mass model, including our fitting procedure and stimulation regime. Finally, we explain the estimation of maximal Lyapunov exponents and construction of Poincaré maps to assess chaoticity.

### 4.1 Empirical Mouse Neural Data

#### 4.1.1 Structural connectivity data

We used tracer-based SC from the Allen Institute (Oh et al., 2014) that is made available in the TVB-data package (https://doi.org/10.5281/zenodo.10128131). The mouse connectome consists of 98 regions and the connection between each regional pair is defined as the ratio between projection density and injection density. The connectivity weights were scaled between 0 and 1. More details about the SC can be found in Melozzi et al. (2017).

#### 4.1.2 LFP data pre-processing

We used publicly available Visual Coding – Neuropixels recordings from the Allen Institute (Siegle et al., 2021). This dataset consists of LFP and spike-sorted data from visual cortical and thalamic areas during passive visual stimulation. The full dataset, including metadata and detailed methodology, is available at: https://allensdk.readthedocs.io/en/latest/visual_coding_neuropixels.html.

We chose a session with recordings from the maximum number of overlapping regions with our SC (*n* = 6, Primary visual area, Field CA1, Field CA3, Dentate gyrus, Subiculum, and Midbrain reticular nucleus). Each session consisted of multiple recording epochs, and we used a 30-second ‘spontaneous’ epoch during which no visual stimuli were presented to the mouse. From the six overlapping regions, we extracted LFP data and interpolated to 1 kHz. To remove channels with low activity, we used root-mean-square (RMS) analysis. We calculated the RMS amplitude for each channel, as the square root of the mean squared voltage, and excluded channels below the threshold of 10% of the median RMS across all channels. Finally, we averaged the remaining signals within each region to obtain one representative time series per region.

### 4.2 Simulation with The Virtual Brain

We used the Larter-Breakspear model (Breakspear et al., 2003; Breakspear and Stam, 2005) to simulate brain dynamics. The Larter-Breakspear model is a conductance-based, three-dimensional dynamical system described by the following equations:

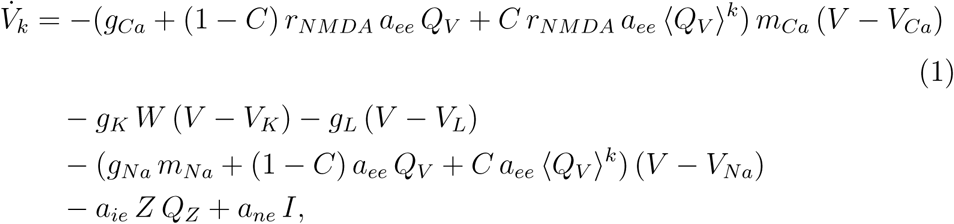

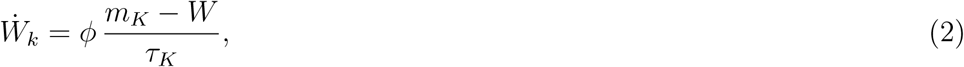

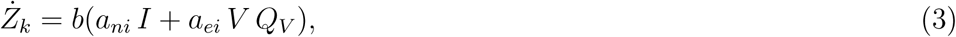

where V denotes the mean membrane potential of the excitatory pyramidal population, Z denotes the mean membrane potential of the inhibitory interneuron population, and W is the proportion of open potassium ion channels.

*Q_V_* and *Q_Z_* are mean firing rates of excitatory and inhibitory populations, respectively, and are defined as

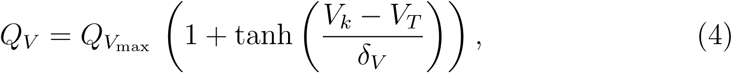

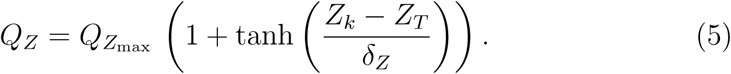

The default parameter values taken from Sanz-Leon et al. (2015) are listed in Table 1.

In TVB, conduction delay between nodes is determined by dividing the estimated tract lengths with a conduction speed. In mice, axonal fibers are less myelinated, shorter, and slower compared to those in human brains, and conduction speeds range between 0.2 mm/ms and 3.5 mm/ms (Wang et al., 2008; Spiegler, 2021). For our simulations, we used a conduction speed of 0.3 mm/ms. Additionally, we explored a broader range of conduction speeds within this range to investigate their impact on network dynamics in 2.3. Note that Allen tracer data does not include information on tract lengths. We used the distance between the centres of the brain regions as a proxy to tract lengths between regions.

#### 4.2.1 Fitting with empirical LFP data

To fit our model to the empirical LFP data from the Allen Institute, we selected overlapping regions between the SC and LFP data, as described in 4.1.2. The fitting target was a 6 × 6 functional connectivity (FC) matrix derived using Pearson correlation on empirical LFP data. For each simulation run, we calculated the FC matrix and reduced it to the same dimensions as the empirical data, by extracting the overlapping regions. The fit between the empirical and simulated FC matrices was evaluated using the root mean squared error (RMSE).

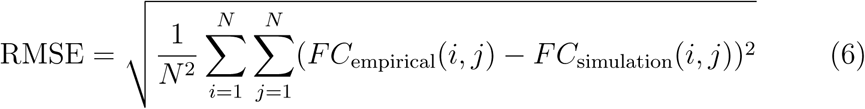

#### 4.2.2 Stochastic Grid Search Optimization

Fitting the Larter-Breakspear network system is challenging due to the high number of parameters and the wide array of possible chaotic regimes. A brute force grid search is impractical for the 31 different model parameters, as even a limited search of 20 values per parameter would result in over 2 · 10^40^ possible combinations. Shortening the simulated time series to speed up computation isn’t viable, as the time series must be long enough to produce accurate FC matrices. To make the problem more manageable, we reduced the parameter space by focusing on key parameters: Nernst potentials, ion channel conductivities, the variance of the excitatory threshold *δ_V_*, and the coupling parameter *C*. These parameters are known to significantly influence the dynamics, as shown in previous studies (Breakspear et al., 2003; Antal et al., 2023). The parameters and the ranges of values used for fitting are shown in Table 1. To further increase fitting speed, we employed stochastic grid search optimization, combining random and grid search methods, commonly used in machine learning hyperparameter tuning. This approach reduces computational costs while maintaining a high likelihood of finding optimal parameter combinations as the number of iterations increases. We ran multiple stochastic grid searches and selected the best-performing combination based on the lowest RMSE, as defined in equation (6). Computations were performed in parallel on the Berlin Institute of Health High-Performance Computing Cluster (BIH HPC). To further refine the fit, we repeated the procedure with a smaller localized parameter grid around the initially found optimum. The evolution of the RMSE fitting metric across iterations is shown in Supplementary files (Fig. S1).

#### 4.2.3 Implementation of External Stimulation

We delivered external stimuli to our simulations to examine the system’s response. Stimuli were delivered to the unitless state variable V (excitatory current, 1) of right primary motor area with an amplitude of 0.1 and a duration of 50 ms. For all simulations, stimulation onset was at 2000 ms and was delivered either once or repeatedly every 500 ms.

### 4.3 Analysis of Chaos in Brain Network Models

#### 4.3.1 Maximal Lyapunov Exponents

A system with sensitive dependence on initial conditions, as described by Strogatz (1994), has neighboring orbits that separate exponentially fast, indicated by a positive Lyapunov exponent. For a one-dimensional map, starting with an initial condition *x*_0_ and a nearby point *x*_0_ + *δ*_0_, the separation *δ_n_* after *n* iterations follows |*δ_n_*| ≈ |*δ*_0_|*e^nλ^*, where *λ* is the Lyapunov exponent. A positive *λ* signals chaotic dynamics. In an *n*-dimensional system, there are *n* Lyapunov exponents, forming the Lyapunov spectrum {*λ*_1_*, …, λ_n_*}, with the maximal Lyapunov exponent (MLE) indicating the long-term separation rate. While some systems allow analytical expressions for Lyapunov exponents, more complex systems require numerical methods to estimate them. Algo-rithms by Rosenstein et al. (1993) and Eckmann et al. (1986) estimate the Lyapunov spectrum or MLE from time series data. We used the algorithm from Rosenstein et al. (1993), implemented in the nolds Python package (Schölzel, 2020), to derive the MLE.

#### 4.3.2 Poincaré Map Analysis

To gain a better understanding of the underlying chaotic dynamics of a system, it is often useful to find a suitable transformation of the phase space dynamics on a lower dimensional manifold. The Poincaré map is an embedding of the dynamical system on a lower dimensional subspace and is used to indicate its chaoticity.

To construct Poincaré maps from time series data, we use a planar cut through the 3D phase space by fixing the state variable Z, which represents the membrane potential of the inhibitory population. We recorded the order of the crossings only from below to effectively reduce the 3D dynamics to two dimensions. In a deterministic, non-chaotic system with stable closed loops, the Poincaré map collapses to a single point, indicating that every subsequent crossing maps onto the same point. An *n*-cycle, on the other hand, appears as *n* discrete points, directly reflecting the number of cycles. To eliminate initial transient dynamics, we excluded the first 1000 ms of simulate data. We then analyzed the sequence of crossing indices using a heat map.

It is also possible to reduce the dimensionality of the phase space by mapping the consecutive local maxima (or minima) of the time series onto each other. To do this, we derived an algorithm that localizes the local maxima and plots them in a (*V* (*n*+1)*, V* (*n*)) space, thus applying the same principle as the Poincaré map. A key benefit of this method is that it removes the need to fine-tune an appropriate intersection plane. To distinguish between chaotic and non-chaotic deterministic dynamics, we employed the local maximum method, which offers a robust criterion for identifying underlying dynamic regimes. This method was applied to both chaotic and non-chaotic dynamical regimes of the Lorenz system (Lorenz, 1963), demonstrating its capacity to differentiate between these regimes, as illustrated in Fig. S4. Additionally, we examined a non-chaotic system subjected to weak additive Gaussian white noise, shown in Fig. S3, to assess the method’s sensitivity to stochastic perturbations. While the local maximum method effectively discriminates between deterministic regimes, care must be taken when interpreting results in noisy systems, as stochastic fluctuations can generate patterns resembling chaotic behavior and may thus confound classification. In the present study, however, all simulations were conducted in the absence of stochastic noise, ensuring that the local maximum method remains a reliable tool for distinguishing between chaotic and non-chaotic dynamics.

## 5 Acknowledgements

PR acknowledges support by EU Horizon Europe program Horizon BRIDGE (101219311), EBRAINS2.0 (101147319), Virtual Brain Twin (101137289), EBRAINS-PREP 101079717, AISN 101057655, EBRAIN-Health 101058516, EIC grant PHRASE 101058240, by the Digital Europe Programme TEF-Health (101100700), SHAIPED (101195135), CoordinaTEF (101168074) German Research Foundation SFB 1436 (project ID 425899996); SFB 1315 (project ID 327654276); SFB 936 (project ID 178316478; SFB-TRR 295 (project ID 424778381); SPP Computational Connectomics RI 2073/6-1, RI 2073/10-2, RI 2073/9-1; DFG Clinical Research Group BECAUSE-Y 504745852, Berlin University Alliance OpenMake, the Virtual Research Environment at the Charité Berlin and EBRAINS Health Data Cloud and the Berlin Institute of Health and Foundation Charité.

## Declaration of generative AI and AI-assisted technologies in the manuscript preparation process

During the preparation of this work the authors used ChatGPT (chatgpt.com/) in order to improve the semantics of already written text. After using this tool/service, the authors reviewed and edited the content as needed and take full responsibility for the content of the published article.

