## Supplementary for "From Ions to Chaos: exploring a whole-brain modelling framework in the mouse"

### Supplementary Materials

#### 1 Stochastic Grid Search Optimization

A stochastic grid search method was employed to identify the optimal parameter set for the 9 selected parameters. The fitting metric used was the Root Mean Square Error (RMSE). Multiple fitting loops were executed in parallel on the High Performance Cluster (HPC) at the Berlin Institute of Health (BIH). This approach allowed us to efficiently explore the parameter space and converge on the best-fitting parameters.

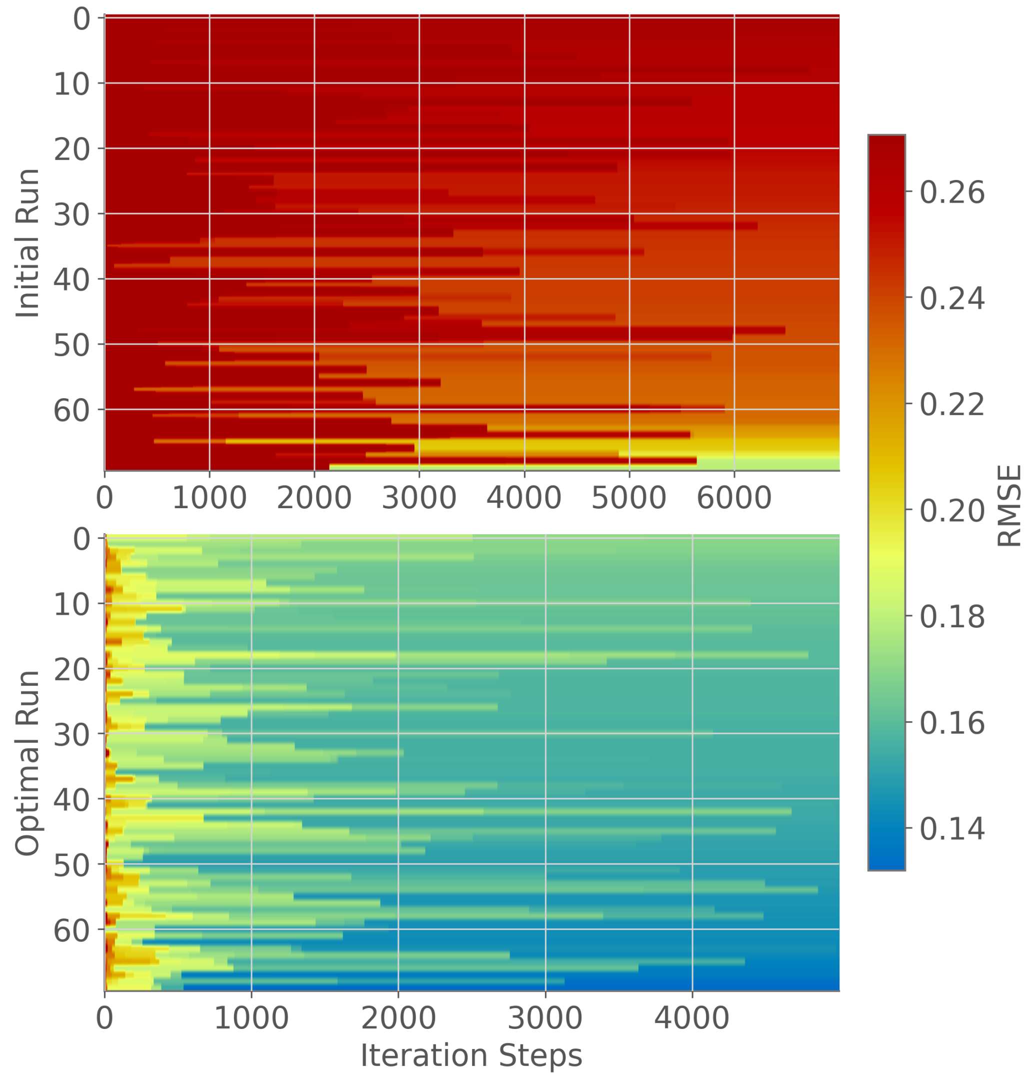


Figure 1: Plot showing the RMSE metric of iterations for a localized run around the initial parameter exploration (top) and finer search around the optimum (bottom). The plot demonstrates the convergence of the fitting process, with the RMSE decreasing over successive iterations, indicating improved parameter fitting. The heatmap displays the best fitting metric at the end of iterations for each parallel run. The parallel localized fitting run (top) converges to a lower RMSE value compared to the initial parameter exploration (bottom) which is displayed with the heatmap. Several initial runs like in the plot (bottom) were performed to localize the area of minima, guiding subsequent fine-tuning runs.

#### 2 Chaotic Dynamics

The study of chaotic dynamics is a field that derived from classical dynamical systems theory. The Lorenz system described in Lorenz (1963) is an atmospheric system governed by 3 coupled non-linear ordinary differential equations.

$$\dot{x} = \sigma(y - x), \tag{1}$$

$$\dot{y} = x(\rho - z) - y, \tag{2}$$

$$\dot{z} = xy - \beta z, \tag{3}$$

$x, y, z$ are the state variables of the system. The variables $\dot{x}$, $\dot{y}$, and $\dot{z}$ represent the derivatives with respect to time $t$. The parameters $\sigma$, $\rho$, and $\beta$ are system parameters governing the behavior of the system. For the parameter combination $\sigma = 10.0, \rho = 28.0, \beta = \frac{8}{3}$, the Lorenz system is chaotic, and the dynamics create the butterfly attractor. The system is unstable arbitrary initial conditions, and the trajectories exponentially diverge from each other. In Fig. 2 we see the chaotic trajectory of the Lorenz system. The trajectory never overlaps but get arbitrarily close to each previous point in the phase space. Even though the dynamics of the system are unstable, the system is bounded in the phase space to an attractor.

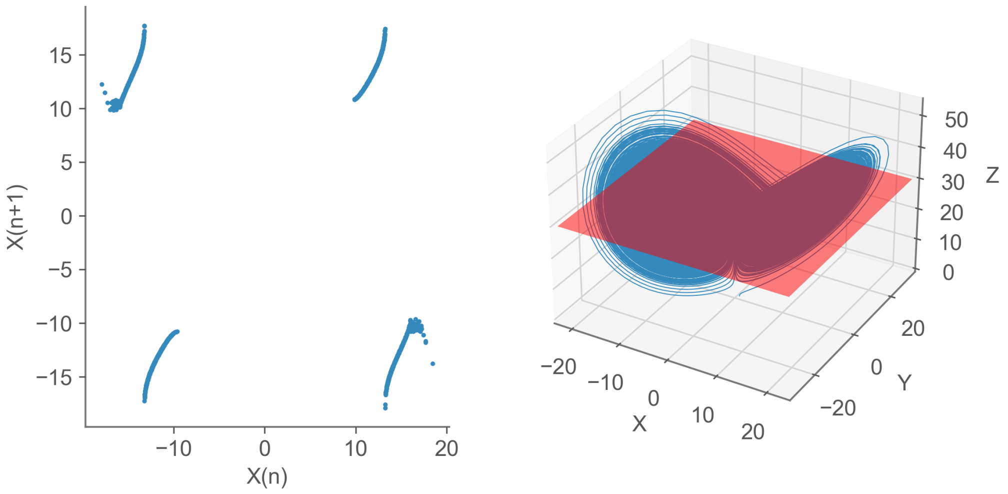


Figure 2: (Right) Three dimensional trajectory of the Lorenz system in the chaotic regime. The trajectory is non-overlapping and creates the strange attractor of the Lorenz system over time. Poincaré section indicated as red plane. (Left) Resulting Poincaré map from plane crossing method that captures consecutive Poincaré points on $x(n), x(n+1)$ axes for discrete and finite amount of $n \in \{1, ..., N\}$. Only crossing from below to above the plane was registered as valid Poincaré point.

### 3 Poincaré Maps

#### 3.1 Plane Crossing Method

The Poincaré map for the Lorenz system (Lorenz, 1963) in the chaotic regime was derived by fixing a $Z$-plane at $Z = 30$ and recording all crossings from below to above the plane. This projection reduces the 3D dynamics to a 2D subspace, plotting the first return points on the $x(n), x(n+1)$ plane. This can be seen in Fig. 2. In chaotic systems, the number of points on the Poincaré map grows indefinitely with simulation time due to the non-intersecting nature of chaotic trajectories. In contrast, non-chaotic systems produce a finite number of return points, corresponding to the number of cycles and the chosen hyperplane.

#### 3.2 Local Maximum Method

We construct a simple example of a use case for the local maximum method for constructing Poincaré maps.

We simulate the following time series.

$$x(t) = \sin(t) + \cos(2.2t) + 0.01 \cdot S,$$

where $t \in (0, 40\pi)$ and $S \sim N(0,1)$ marks a weak additive gaussian white noise term.

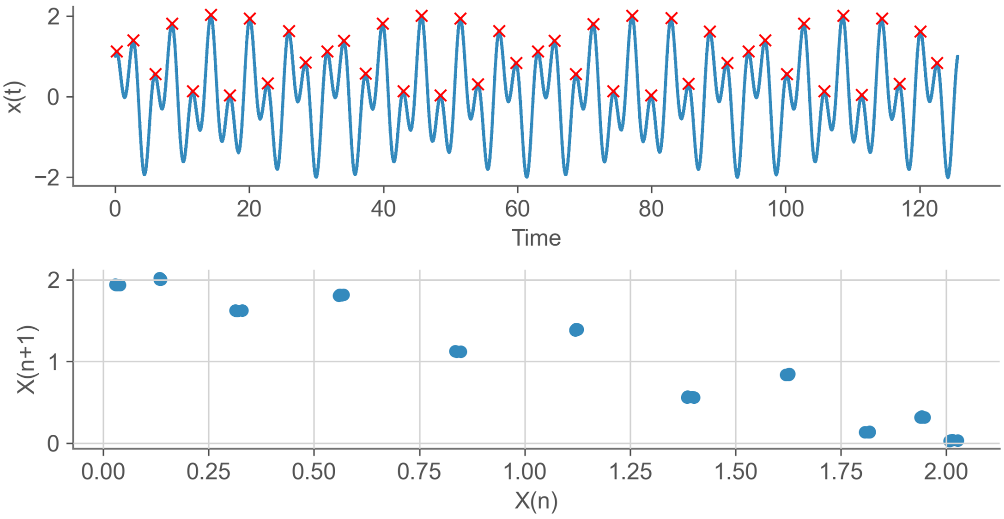


Figure 3: Time series of an oscillatory signal with stochastic noise. Poincaré map derived with local maximum approach.

Fig. 3 shows the time series of $x(t)$ and the resulting Poincaré map. The dynamics of the time series can be described as a $n$-cycle of order $n = 11$, so the first and $12th$ local maximum should be identical. Because of weak additive Gaussian white noise we will not see exact equality. Nevertheless, the Poincaré map clearly distinguishes 11 different point clouds that are aligned in a shape of an ellipse. If we apply the local maximum approach to the Lorenz system we will see that the resulting Poincaré map also shows a similar complex geometric structure as before. The Poincaré map is constructed by tracking the local maxima of the $x$-Variable of the Lorenz system.

In Fig. 4 in panels (A),(B) we see how the resulting Poincaré map has 4 different wings to it, that are characterized by a high number of points. The chaoticity of the dynamics can be directly deduced from the geometry of the Poincaré map. As the length of the analyzed time series increases the number of points in the Poincaré map grows continously.

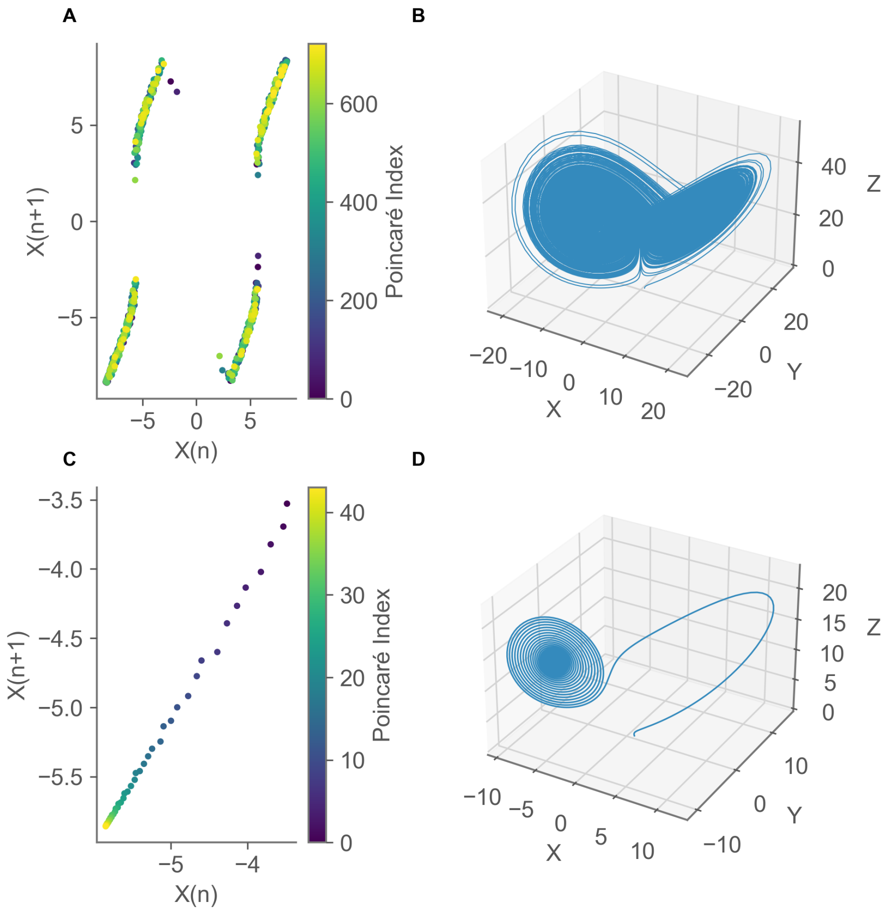


Figure 4: Time series of the chaotic Lorenz system and Poincaré map derived with local maximum approach. (A), (B) chaotic Lorenz system. (C), (D) non-chaotic converging Lorenz system.

Lastly, we want to showcase what the Poincaré map of a deterministic non-chaotic dynamical system would look like. We can make use of the

Lorenz system and alter the parameters to $\sigma = 10.0, \rho = 14.0, \beta = \frac{8}{3}$, where we decreased $\rho$ and therefore stabilize the system. The Lorenz system is still oscillating with these new parameters but also converges towards a stable fixed point.

In Fig. 4 in panels (C), (D) we see the resulting Poincaré map of the stable oscillatory Lorenz system. The trajectory in the 3D-phase space is stable and converges to a fixed point. This convergence in the $(x, y, z)$ phase space is directly translated into a convergence on the Poincaré map. On the Poincaré map we can easily see how the consecutive local maxima points converge towards a fixed point where, which indicates that the underlying higher dimensional dynamics are in fact not chaotic.

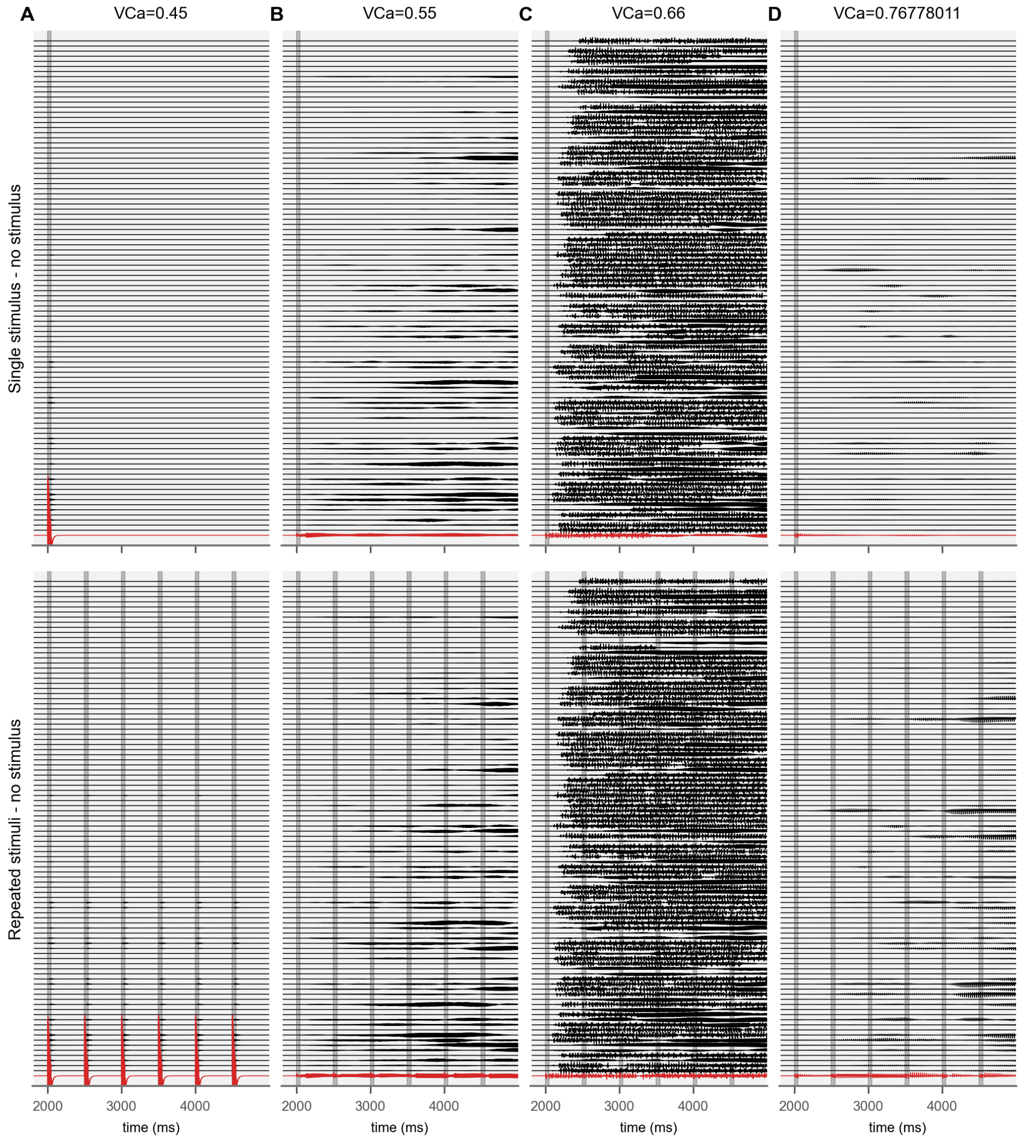


Figure 5: Calcium channel activity modulates the simulated system's sensitivity to external stimuli. The difference between stimulus and no stimulus conditions from simulations using identical initial conditions are plotted, thereby demonstrating the effect of stimulus. All 98 regions of the Allen mouse connectome (Oh et al., 2014) are plotted. Stimuli were delivered to right primary motor area, plotted in red. Other regions are plotted in the order of connectivity strength to the stimulated region, from bottom (strongest connectivity) to top (weakest connectivity).
